# *Neoehrlichia mikurensis*, where art thou? A negative analysis of 87 Swedish wild boars (*Sus scrofa*)

**DOI:** 10.64898/2026.09.29.755505

**Authors:** Hanna Lindell, Andrew Jenkins, Kayla C. Evens, Kari Lehtilä, Patrik Dinnétz

## Abstract

*Neoehrlichia mikurensis* is widely detected in tick populations across Europe, yet its host range beyond small rodents remains poorly understood. Large wild mammals are frequently exposed to tick vectors, but their role in the transmission cycle of this bacterium is unclear. Wild boars (*Sus scrofa*) are among the most abundant large mammals in southern Sweden and commonly inhabit tick-infested environments, making them a potentially relevant host species.

In this study, spleen samples from 87 wild boars collected between 2023 and 2025 across southern and central Sweden were analysed for the presence of *N. mikurensis* DNA using quantitative polymerase chain reaction. *N. mikurensis* DNA was not detected in any of the analysed samples, despite sampling in regions where the bacterium is known to be prevalent in ticks. These results indicate that wild boars may have a limited role in the maintenance of N. *mikurensis* in Sweden, although the possibility of rare or transient infection cannot be excluded. Despite being based on negative findings, this study contributes to a more comprehensive understanding of the host range of *N. mikurensis* and highlights the importance of continued evaluation of wildlife species in the transmission dynamics of tick-borne pathogens.

## Introduction

*Neoehrlichia mikurensis* is an emerging tick-borne pathogen within the family *Anaplasmataceae*. This gram-negative, intracellular bacterium is widely distributed in nature and has been detected throughout a large part of Europe [1]. In humans, *N. mikurensis* can cause severe symptoms, including fever and myalgia in immunocompromised individuals. However, the detection of *N. mikurensis* in blood donors [2,3] suggests, however, that the bacterium may also be capable of establishing chronic, subclinical infections in immunocompetent individuals. A distinguishing feature of the bacterium is its strong tropism for vascular endothelial cells which can cause thromboembolic complications such as vasculitis and pulmonary embolism [4,5].

*N. mikurensis* has been detected in several tick species, but *Ixodes* spp. are considered the primary vectors [6]. The prevalence of *N. mikurensis* in questing *Ixodes ricinus* ticks in Europe varies between regions, ranging from 0.1% in Denmark to 24.3% in Hungary. In southern Sweden, a prevalence of approximately 6.0% has been reported in questing ticks, indicating a substantial presence in the local tick population [7,8].

Small rodents, particularly species of the genera *Apodemus, Microtus* and *Myodes*, are considered the main natural reservoir hosts [6]. However, *N. mikurensis* can infect humans and also dogs [6,9], which raises the question of whether other wild or domestic animals may contribute to its enzootic cycles, and a recent conference report [10] describing findings in European badgers (*Meles meles*), Brown bears (*Ursus arctos arctos*), chamois (*Rupicapra rupicapra*), mouflons (*Ovis gmelini musimon*), and wild boars (S*us scrofa*) in Croatia suggest that might be the case. Engorged ticks collected from wild boars in the Netherlands and\Belgium have also tested positive for *N. mikurensis* [11]. Thus, the host competence of wild boar deserves further attention.

The aim of this study is to examine the relevance of the newly established large mammal species, the wild boar, as a potential host of *N. mikurensis* in Southern Sweden.

Understanding the full host range of *N. mikurensis* is important, as it may influence transmission dynamics, geographic spread, and risk of human exposure. Wildlife species such as wild boars, which are increasingly abundant across Europe and frequently inhabit areas where human and wildlife habitats intersect, are particularly relevant in this context [12,13] .

## Materials and methods

### Study area and sample collection

In February 2023, 229 local hunting associations (jaktvårdskretsar) across Sweden were contacted regarding participation in a study focusing on tick-borne pathogens in wild boar. The hunters contacted were active in areas stretching from the southern border of Sweden (Skåne) up to Dalarna, as wild boars are sparse further north. Of those contacted, 32 responded positively and agreed to take part in the study.

Between March 2023 and March 2025, participating hunters collected spleen samples from wild boars harvested for human consumption during field dressing. Immediately after collection, approximately 1 × 1 cm of spleen tissue from each animal was preserved in 70%ethanol and sent by post to Södertörn University. Upon arrival, samples were stored at −80 °C until laboratory processing.

In total, samples from 89 wild boars were received, of which 87 spleen samples were analyzed for the presence of *N. mikurensis*. Previously, spleen has successfully been used to detect N. mikurensis in rodents [14]. The geographical distribution of the tested spleen samples was as follows (number of wild boars in parentheses): Halland (*n* = 1), Kronoberg (6), Örebro (3), Östergötland (2), Skåne (4), Södermanland (32), Stockholm (2), Uppsala (12), Västmanland (3), and Västra Götaland (22) (Figure 1).

**Figure 1.**
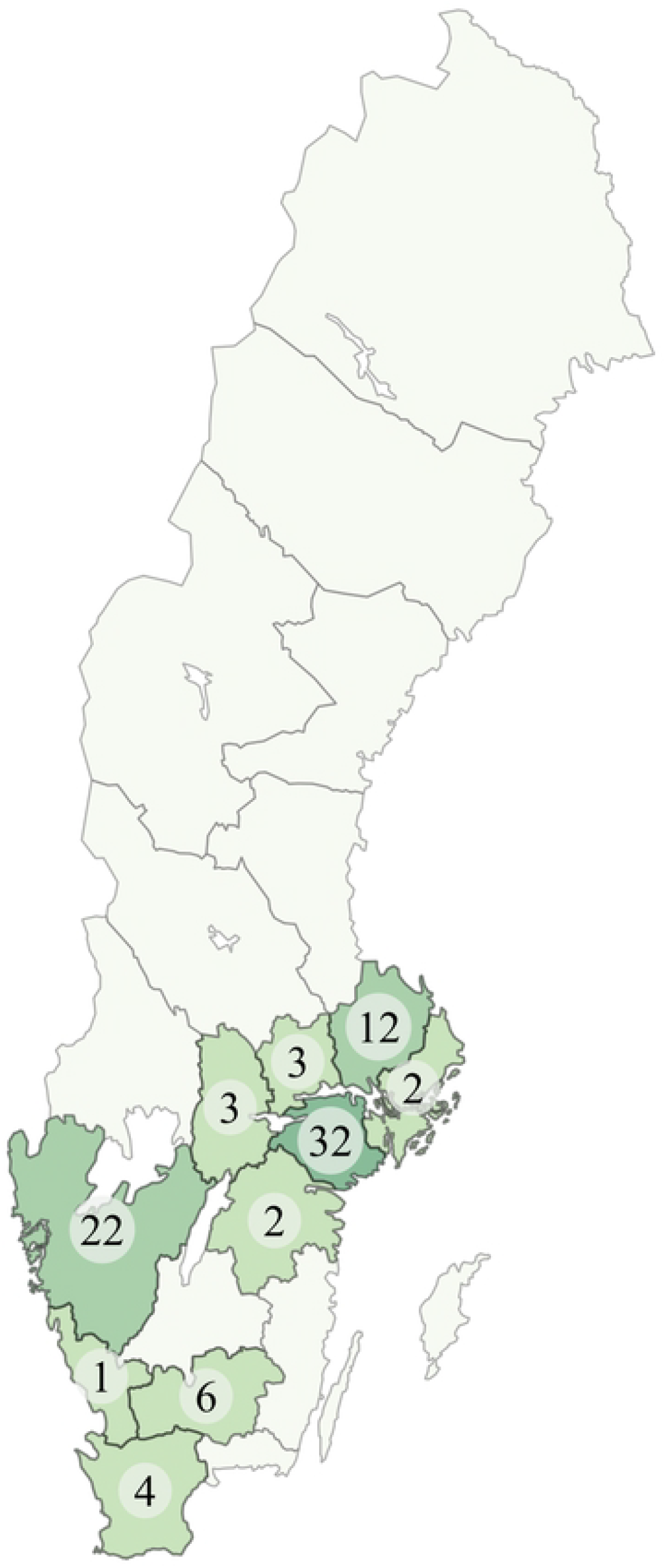
Geographical distribution of wild boar spleen samples collected for *N. mikurensis* testing. Numbers indicate the number of samples per county; darker shading represents higher sample counts. Administrative boundaries are based on the Lantmäteriet geodata product *Kommun, län och rike* (© Lantmäteriet, CC BY 4.0). The data were processed and visualized by the authors using QGIS

In addition, ticks found on the animals during field dressing were opportunistically collected by hunters. In total, 30 ticks were obtained from 17 of the 89 sampled wild boars, distributed across 10 counties. The majority were adult females (n = 26), while the remainder were nymphs (n = 4). Given the limited number of ticks and their wide geographical dispersion, testing for *N. mikurensis* was considered unlikely to yield robust or representative estimates of local N. mikurensis prevalence, and these ticks were therefore not analyzed in the present study.

The geographical distribution of the sampled wild boars largely reflects regions with high wild boar density in Sweden as estimated from wild boar-related traffic accidents, which is commonly used as a proxy for population density[15].

Ethical approval was not required for this study, as it did not involve live animals or animals killed specifically for scientific purposes. All wild boars included in the study were legally hunted for human consumption, in accordance with the Swedish Hunting Act (SFS 1987:259).

### Detection of *Neoehrlichia mikurensis*

DNA extraction was performed at Södertörn University, Huddinge, Sweden, using the QIAamp Fast DNA Tissue Kit (Qiagen), according to the manufacturer’s protocol.

. The resulting DNA eluate was subsequently sent to the University of Southeastern Norway, Bø, Norway, for quantitative PCR (qPCR) analysis.

Quantitative PCR targeting a 131 bp conserved segment of the *groEL* gene of *N*.*mikurensis* was performed as previously described [16]. A 10-fold serial dilution of a synthetic positive control plasmid pNeo [13] from 1.6 × 10^6^ to 16 copies was included in each run and served as a control on sensitivity. Dissociation analysis was performed at the end of the run to determine the melting temperature of amplicons. A detectable amplification curve and a melting temperature between 71.7 and 75 °C was the working definition of a positive sample.

Spiking analysis was used to control for inhibition as previously described [17] Briefly, PCR reactions were set up containing sample DNA and 1.6 x 10^4^ of positive control plasmid, pNeo. Samples were considered inhibitory if the presence of their DNA resulted in a right-shifted amplification curve (increased Cq) or if no amplification curve was detected. Where inhibition was detected, the sample was diluted 1:5 or 1:10 in molecular biology grade water and PCR was repeated, in parallel with a spiked, diluted sample to confirm that inhibition was successfully eliminated.

## Results and Discussion

Although the pathogens established presence in Swedish tick populations and its detection in other wildlife hosts across Europe[7,9,10], *Neoehrlichia mikurensis* DNA was not detected in any of the 87 wild boar spleen samples analyzed in this study (Table 1).

**Table 1.** Summary of detection of *N. mikurensis* in wild boar samples by gender and age group.

|  | Number | Percentage of<br>total<br>sample numbers | Number of Positives |
| --- | --- | --- | --- |
| **Gender** |  |  |  |
| Male | 45 | 51.7 | 0 |
| Female | 42 | 48.3 | 0 |
| Total | 87 | 100 | 0 |
| **Age** |  |  |  |
| Piglet: < 4 months | 0 | 0 | 0 |
| Juvenile: 4-12 months | 46 | 52.9 | 0 |
| Adult: >12 months | 40 | 46 | 0 |
| Unknown | 1 | 1.1 | 0 |
| Total | 87 | 100 | 0 |

One previous study has reported *N. mikurensis* in wild boars, however, this result originates from an oral presentation reported in a conference proceeding [10] and therefore provides limited methodological detail and making it hard to compare their result with our study. Additionally, the detection of *N. mikurensis* in engorged *Ixodes* ticks collected from wild boars in earlier studies [11] does not necessarily indicate that the present host individual was infected. If the attached ticks were nymphs or adults, they may have acquired the infection from a previous host and retained it transstadially. Alternatively, larvae may acquire the bacterium through co-feeding transmission, where infected and uninfected ticks feed in close proximity and transmit pathogens locally through the host’s skin without requiring a systemic infection in the vertebrate host. Thus, the presence of *N. mikurensis* in ticks removed from wild boars cannot be used as direct evidence that wild boars themselves are competent or even susceptible hosts. Taken together, these observations indicate that the earlier findings do not provide reliable evidence of infection in wild boars. Nevertheless, they underline the importance of assessing whether wild boars fulfil the conditions required to function as reservoir hosts, especially in light of the discrepancy between infected ticks in the environment and the absence of detectable infection in our material.

To evaluate the significance of a host species in the natural transmission cycle of a tick-borne pathogen, three key factors must be considered: (1) the species population density, (2) the average tick burden per individual, and (3) the reservoir competence, defined as the proportion of ticks that acquire the infection after feeding on an infected host. An individual species’ ability to act as a reservoir host is determined by both its biology and the characteristics of the pathogen [18].

Applying these criteria to wild boars, the first factor, host abundance, is clearly met as wild boars are both numerous and widely distributed in Sweden and large parts of Europe [12,13] . However, the second factor, average tick burden per individual, is less clear. While wild boars do host *Ixodes* spp. ticks, the intensity of infestation appears to be considerably lower compared to other large wild mammals. In the present study, hunters were asked to check the animals for attached ticks at the time of sampling and found a total of 30 ticks from 17 wild boars. However, the collection was not mandatory and was dependent on practical circumstances such as time constraints during field dressing. Consequently, the number of ticks recorded is likely an underestimation of the true tick burden. These observations should therefore be interpreted cautiously and cannot be used to quantitatively assess infestation intensity.

In contrast, Fabri et al. (2021) conducted systematic tick collection and found that wild boars had the lowest tick infestation rates among five ungulate species in south-central Sweden: fallow deer (*Dama dama*), moose (*Alces alces*), red deer (*Cervus elaphus*), roe deer (*Capreolus capreolus*), and wild boar (*Sus scrofa*). This applied across all life stages of *I. ricinus*. For larvae, wild boars showed 0% infestation prevalence, compared to 18% in fallow deer and 10% in roe deer. For nymphs, wild boars had an infestation prevalence of 17%, while fallow deer, roe deer, red deer, and moose, had rates of 97%, 94%, 66% and 39%, respectively. Adult female ticks were found on only 3% of wild boars, in contrast to 95–98% in the deer species. These findings suggest that wild boars are less infested by ticks in general. However, in the same study, *Anaplasma phagocytophilum* was detected in 24 out of 34 wild boars tested, indicating that they must have been exposed to ticks at some point during their lifetime [19]. This discrepancy highlights the uncertainty regarding tick infestation rates in wild boars and indicates that occasional or transient tick infestations could still be epidemiologically relevant, emphasizing the need to further investigate the role of wild boars in tick-borne pathogen transmission cycles.

Wild boars’ reservoir competence for *N. mikurensis* is still unknown. Rodents have been identified as the main reservoir hosts for *N. mikurensis* both in field studies showing higher prevalence in feeding ticks collected from rodents compared to questing ticks, as well as in xenodiagnosis experiments, where uninfected ticks feed on infected hosts to test for transmission [14,20,21] However, to our knowledge, such experiments have not been conducted on any other species than rodents.

Since our findings are entirely negative and seem to contradict those of a previous report [10], the possibility of our results being a negative artefact needs to be considered. The possible causes of false negative results are inhibition, incorrect choice of sample, too few samples, sample degradation and unsuitable PCR test.

Thorough procedures to control and exclude inhibition were implemented and inhibition can be excluded as a source of false negative results.

Our choice to analyse the spleen was based on an earlier study concluding that the spleen and kidneys were the most effective organs for detecting *N. mikurensis* in rodents [14] . Therefore, we assumed that the spleen should also be a suitable organ for pathogen detection in wild boars. Although 87 individuals represent a relatively large sample size in wildlife research, it is difficult to determine what constitutes a sufficient sample size when investigating pathogens with potentially low prevalence. Future studies could benefit from larger sample sizes, repeated sampling over time, and inclusion of multiple tissue types to increase the likelihood of detecting transient or low-level infections.

Host-specific internal controls were not included to assess DNA integrity in individual samples. However DNA extracts from the present study have been successfully used in parallel molecular analyses as part of a larger project, indicating that amplifiable DNA was recovered from the samples (Lindell et al, unpublished).

The PCR test used in this study has been validated for analytical sensitivity according to the MIQE guidelines [16] and has also been successfully used to detect *N. mikurensis* in ticks [16,22,23] and humans [24] Sensitivity controls were included in all PCR assays and analytical sensitivities of 16 copies were routinely obtained. False negative results due to assay inadequacy thus seem unlikely.

In conclusion, given the relatively high prevalence of *Neoehrlichia mikurensis* in ticks (6% in Sweden) and the absence of infection in wild boars despite their relatively frequent exposure to ticks, our findings suggest that wild boars may not be susceptible to chronic infection.

Furthermore, if infections were acute and severe, clinical signs or elevated mortality would probably have been observed, particularly given the high abundance and visibility of wild boars in the landscape. The lack of such observations supports the notion that *N. mikurensis* either does not infect wild boars or does so in a transient, mild, or subclinical manner.

Despite the negative findings, this study contributes valuable data to the limited body of research on *N. mikurensis* in wildlife hosts beyond rodents. Wild boars are epidemiologically relevant due to their abundance, expanding distribution across Sweden and Europe, and frequent exposure to tick vectors. Their wide-ranging behavior and overlap with human environments further reinforce their potential role in the ecology of tick-borne pathogens. Future studies should aim to clarify the full host range and identify environmental drivers of *N. mikurensis* transmission across different ecological settings, including systematic collection and testing of ticks from wild boars to better assess local pathogen exposure and circulation.

## Data Availability

All relevant data are provided within the manuscript and supplementary information files.

## Acknowledgments

The authors sincerely thank the hunters who participated in this study. Their generous collaboration and contribution of samples were essential, and this research would not have been possible without their support.

## Supporting information

S1 Appendix: qPCR-result report, amplification plots, standard curves and melt curves.

